# Mitigating Proinflammatory SASP and DAMP with Urolithin A: A Novel Senomorphic Strategy

**DOI:** 10.1101/2025.01.07.631588

**Authors:** Anna Barkovskaya, Ashley Brauning, Manish Chamoli, Anand Rane, Julie K Andersen, Amit Sharma

## Abstract

Senescent cells are known to contribute to aging and age-related diseases. One key way they influence aging is by secreting senescence-associated secretory phenotype (SASP) factors along with several damage-associated molecular pattern (DAMP) molecules. Consequently, inhibiting SASP and DAMP signaling (senomorphics) has emerged as a therapeutic strategy. Urolithin A (UA), a gut-derived metabolite produced from ellagitannins and ellagic acid found in berries, nuts, and pomegranates, has demonstrated potent anti-inflammatory properties and protective effects against aging and age-related conditions in experimental models. Here we demonstrate that UA lowers the expression and release of pro-inflammatory SASP and DAMP factors at least in part, on the downregulation of cytosolic DNA release and subsequent decrease in cGAS-STING signaling.

## Introduction

Aging is associated with increased systemic sterile inflammation (inflammaging), which promotes several age-associated diseases (Franceschi and Campisi, 2014). Key drivers of inflammaging include senescence-associated secretory phenotype (SASP) factors released by senescent cells (Olivieri et al., 2018). While the exact components of SASP vary between different senescent cells and tissues, core SASP factors include pro-inflammatory chemokines, matrix-degrading enzymes, and several damage-associated molecular pattern (DAMP) molecules (Coppe et al., 2010). Pharmacological inhibition of SASP using small molecules called senomorphics has been proposed as a potential intervention for age-associated disease (Kim and Kim, 2019; Lagoumtzi and Chondrogianni, 2021). However, such treatments include several flavonoid inhibitors of the p38 MAPK/NF-κB pathway (Freund et al., 2011; Zhang et al., 2023), free radical scavengers, and Janus kinase (JAK) pathway inhibitors (Niedernhofer and Robbins, 2018) that are non-selective and broadly inhibit pathways also involved in homeostatic immune responses to various physiological challenges, thus limiting their systemic therapeutic application (Zhang et al., 2017).

Here we present data indicating that the gut metabolite Urolithin A (UA) acts as a senomorphic compound. The digestive tract bacteria naturally produce UA through the metabolism of ellagitannins and ellagic acid, which are abundant in berries, nuts, and pomegranates (D’Amico et al., 2021). UA has been reported to be a potent anti-inflammatory agent, alleviating several age-related conditions *in vivo* (D’Amico et al., 2021; Ishimoto et al., 2011; Larrosa et al., 2010). Preclinical studies have also shown its protective role against aging and age-related conditions affecting the muscles, brain, joints, and other organs (D’Amico et al., 2021). In a recent clinical trial, UA supplementation improved muscular endurance in older adults (Liu et al., 2022).

## Results and discussion

To test the effect of UA on senescent cells, we induced senescence in the human fetal lung fibroblast IMR-90 line via treatment with 300 nM doxorubicin (S). We observed robust senescence induction as measured by senescence-associated beta-galactosidase (SA-β-gal) staining (93%) 10 days following doxorubicin treatment compared to non-senescent control controls (<2%) **(Figure 1A A-i,)**. In addition to chemotherapy-induced senescence, we also examined the effects of UA in the context of replicative senescence (RS) (Campisi, 1997). Both doxorubicin-treated S and RS cells displayed significantly elevated expression of the cell cycle checkpoint inhibitors *p16*^*INK4A*^ and *p21*^*CIP*^. Treatment with UA had no significant effect on the expression of these makers **(Figure 1B B-i,)**. Previously, UA pre-treatment has been shown to reduce the percentage of SA-β-gal - positive cells and p53 and p21 expression in senescent Organ of Corti 1 (HEI-OC1) cells and cochlear explants (Cho et al., 2022) or mesenchymal stem cells derived from nucleus pulposus (Cho et al., 2022; Shi et al., 2021). In another study, treatment with UA did not affect the expression of SA-β-gal in human skin fibroblasts undergoing replicative senescence (Liu et al., 2019). While our results show no significant effect on the expression of *p16*^*INK4A*^ and *p21*^*CIP*^ albeit with a trend towards lower expression, it may be due to the diversity of senescence phenotypes.

**Figure 1.**
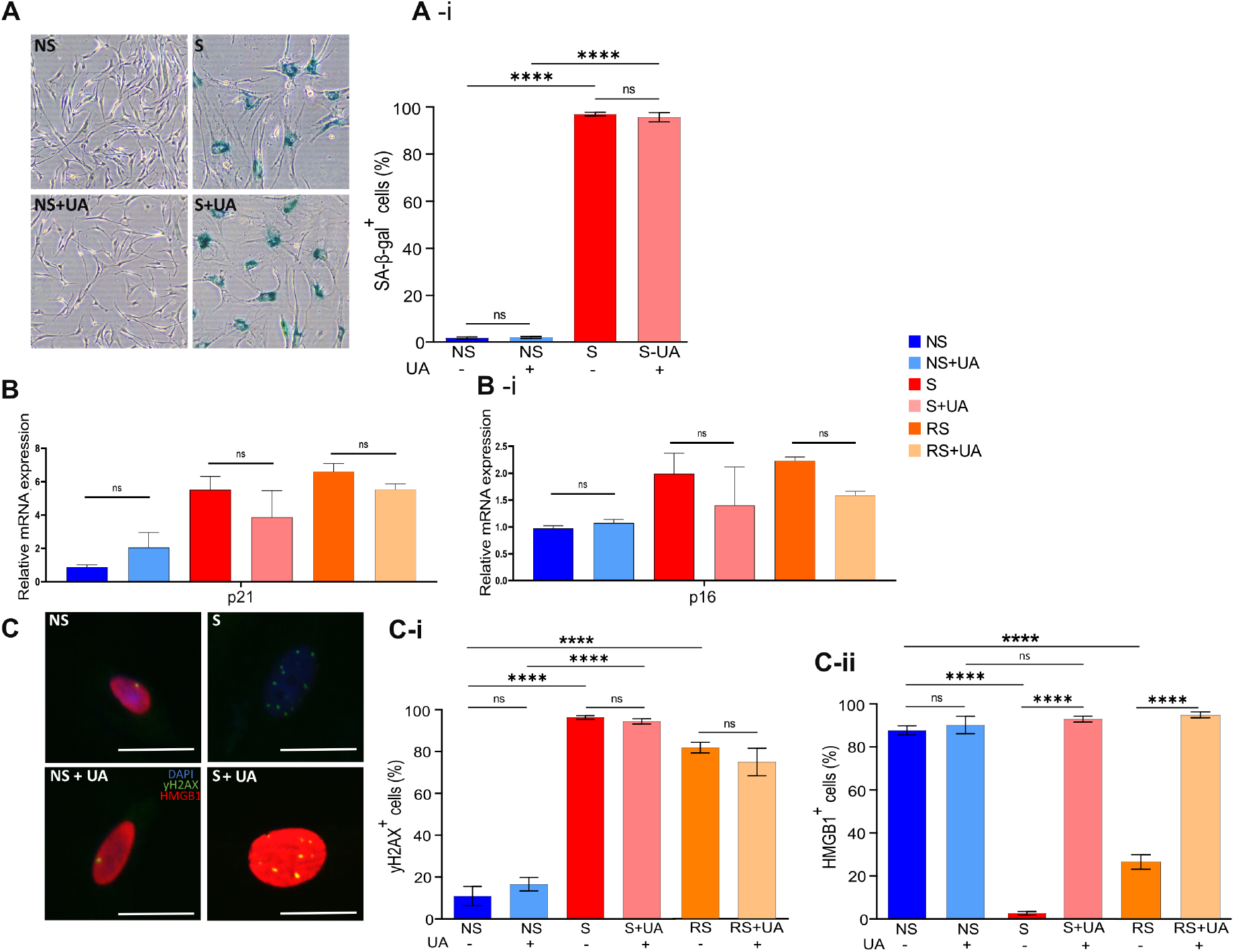
*Doxorubicin treatment induces senescence in human IMR-90 fibroblasts*. **A**.Representative images of SA-B-Gal-stained S and NS with or without the addition of Urolithin A (UA). **A-i**. The proportion of cells staining positive for SA-βGal in NS and S cells with and without the UA treatment. Four fields were quantified per well, n=3. **B, B-i**. Relative p16 (B) and p21 (B-ii) expression in NS, S, and RS cells with and without UA treatment, n=4. **C**. IF staining for γ-H2AX and HMGB1 in NS, S, and RS cells with or without the addition of UA, 10 days post-induction. γ-H2AX green, HMGB1 - red and DAPI - blue. **C-i, C-ii**. The percentage of cells with three or more γ-H2AX foci per cell (C-i) and with nuclear HMGB1(C-ii). All data presented as mean ± SEM. One-way ANOVA. *p <0.033, **p<0.002, ***p<0.0002 ****p<0.0001.

Further confirming senescence induction, most doxorubicin-treated IMR-90 cells had two or more γH2AX foci, indicating double-strand DNA breaks (Hernandez-Segura et al., 2018; Kinner et al., 2008). Treatment with UA did not prevent DNA damage either **(Figure 1C-i)**. The loss in the nuclear localization of high mobility group box 1 (HMGB1), a member of the highly conserved nonhistone DNA-binding high-mobility group protein family, is a well-established senescence marker (Davalos et al., 2013). Its secretion from senescent cells has been reported as part of damage-associated molecular pattern (DAMP) signaling, a key driver of the senescent phenotype (Wiley and Campisi, 2021). Notably, treatment with UA significantly reduced the percentage of cells that lost HMGB1 from the nucleus in both the doxorubicin-treated and RS cells **(Figure 1C, C-ii, Figure 2C-ii)**.

**Figure 2.**
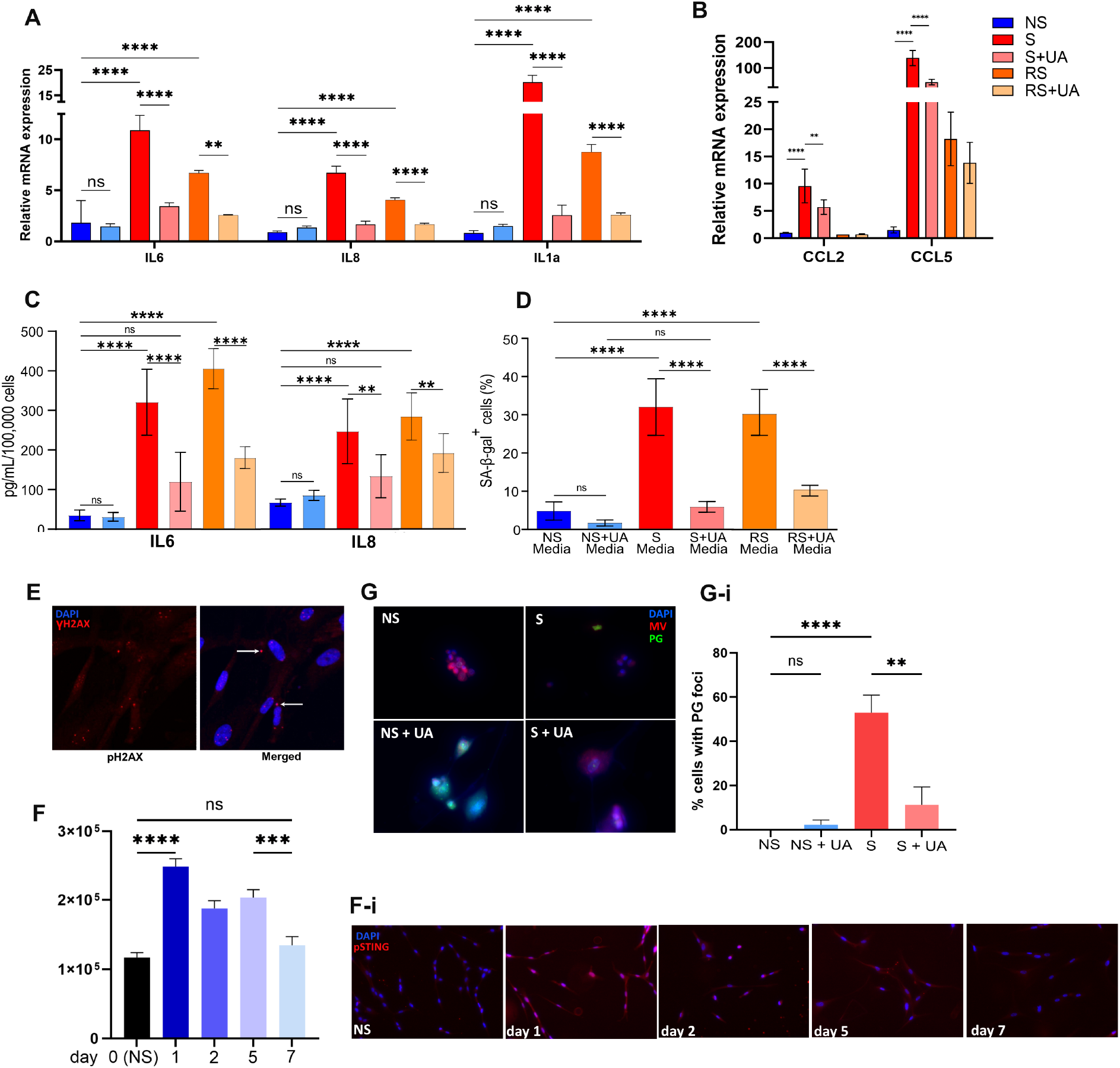
*UA decreases SASP gene and protein expression in senescent cells and down-regulates the c-GAS – STING signaling*. **A**.Relative gene expression of IL6, IL8, and IL1-α in NS, S and RS IMR-90 cells with or without the UA treatment, n=4. **B**. Relative gene expression of CCL2 and CCL5 in NS, S and RS cells with or without UA treatment, n=3. **C**. ELISA on conditioned media collected from NS, S and RS cells with and without UA treatment after 24 hours of incubation. Cytokine concentrations were normalized to 100,000 cells, n=3. **D**. Proportion of SA-βGal-positive cells 6 days after they have been treated with media collected from NS, S or RS cultured with or without UA for 6 days, n=3. **E**. Representative images of the cells labeled with γ-H2AX antibodies and DAPI showing extranuclear DNA in the S cells. **F**. Quantification of the pSTING intensity staining in the NS cells and in cells treated with doxorubicin, evaluated at days 1, 2, 5 and 7, n=3. **F-i**. Representative images of the NS cells and in cells treated with doxorubicin, evaluated at days 1, 2, 5 and 7, labeled for pSTING and DAPI. **G**. Representative images of the NS and S cells with and without the UA treatment, labeled with DAPI, mito-view tracker red and pico-green. **G-i**. Percentage of the cells with extranuclear pico-green foci in the NS and S cells with and without the UA treatment, n=3. All results are presented as a mean and error bars represent SEM. Statistical analysis performed using one way ANOVA. *p <0.033, **p<0.002, ***p<0.0002 ****p<0.0001

Treatment with UA also resulted in a significant reduction in the expression of prototypical SASP factors, including interleukin 6 (*il6*), interleukin 8 (*il8*), and interleukin 1-alpha (*il1α*), in both models of senescence induction. At the same time, UA did not affect SASP gene expression in the non-senescent cells **(Figure 2A)**. Reduced gene expression was followed by a significant decrease in secretion of IL6 and IL8 after UA treatment, as measured by ELISA in both doxorubicin-treated and replicative senescent cells **(Figure 2C C-i,)**.

The primary physiological function of SASP from senescent cells is to recruit immune cells that facilitate tissue repair and regeneration during wound healing. The pro-inflammatory chemokines C-C motif ligand 2 (CCL2) and C-C motif chemokine ligand 5 (CCL5) can recruit and activate macrophages and NK cells, promoting local inflammation (Budamagunta et al., 2021; Luciano-Mateo et al., 2020). As expected, expression of both chemokines was significantly upregulated in senescent cells. Treatment with UA mildly reduced expression, specifically in doxorubicin-treated cells, suggesting that UA has a distinct mechanism of action dependent on the type of senescence induction **(Figure 2B)**.

It has been previously shown that in addition to their pro-inflammatory function, SASP factors propagate senescence to neighboring cells in tissues(Admasu et al., 2023). We, therefore, tested whether UA treatment abrogates paracrine senescence between cells by culturing proliferating IMR-90 in the presence of media collected from either control or UA-treated senescent cells. We found that UA reduced the proportion of SA-β-gal-positive cells in culture, suggesting that it acts as an inhibitor of paracrine senescence **(Figure 2D)**.

Finally, we determined the possible mechanisms associated with UA-mediated cytokine and chemokine regulation. Cellular senescence is associated with the accumulation of cytosolic DNA released from structurally impaired nuclei and mitochondria. Through an evolutionary conserved mechanism of pathogen detection, misplaced DNA fragments are recognized by intracellular DNA sensors, including cyclic GMP-AMP Synthase (cGAS) and the cGAS effector protein Stimulator of Interferon Genes (STING) (Sun et al., 2013). cGAS binding to double-stranded DNA (dsDNA) results in cyclic guanosine monophosphate–adenosine monophosphate (cGMP) buildup as a secondary messenger that binds to and activates STING. STING, in turn, induces the translocation of transcription factors nuclear factor-kappa B (NF-κB) and IFN regulatory factor 3 (IRF3) into the nucleus (Sun et al., 2013). This promotes the expression of pro-inflammatory SASP factors, fueled further by the direct exchange of cGAMP between neighboring cells through cGAMP transporters and GAP junctions, creating a more widespread inflammation (Chen et al., 2016). The cGAS-STING pathway has been proposed as a potential target for pharmacological intervention in disease treatments and controlling inflammation and immunity (Chin et al., 2023). We found that doxorubicin-treated senescent cells indeed showed evidence of cytosolic DNA accumulation as demonstrated by induction of γH2AX foci **(Figure 2E)** and dsDNA-specific pico-green (PG) staining present outside of the nuclear boundaries **(Figure 2G G-i,)**. STING phosphorylation at Ser366 residue was previously shown to be required for STING-dependent activation of IRF3 downstream (Liu et al., 2015). In line with our previous findings, pSTING is significantly more abundant in senescent cells, reaching its highest levels shortly after treatment with doxorubicin before returning to its original level at day 7 **(Figure 2F F-i,)**. We found that the cytosolic DNA foci were significantly reduced following treatment with UA **(Figure 2G-i)**. This suggests that UA down-regulates dsDNA leakage from the nucleus and mitochondria, decreasing the local inflammatory response. These data align with the previous reports that UA enhances cytosolic DNA clearance in cancer cell lines (Madsen et al., 2024). In aged mice, UA supplementation improved muscle strength, reduced inflammation, and promoted autophagic and mitochondrial health (Luan et al., 2021). UA treatment has also been shown to increase self-renewal and prolong T memory stem cell (TSCM) survival, thus preventing T cell exhaustion and promoting effective anti-tumor T cell responses, further supporting its use as an anti-aging therapeutic (Luan et al., 2021). Senescent cells are known to have dysfunctional mitochondria, which results in oxidative stress. Over time, damaged mitochondria can release mtDNA into the cytosol. Cytosolic mtDNA can act as a DAMP molecule, triggering inflammation via pathways such as the cGAS-STING and secretion of the SASP and DAMP. UA is known to activate mitophagy, so UA treatment may reduce cytosolic DNA by activating mitophagy in senescent cells.

Additionally, while cGAS-STING activation is primarily an inflammatory response, it can act as a feedback mechanism to promote mitochondrial quality control by activating mitophagy. Studies have demonstrated that cGAS-STING activation can upregulate mitophagy-related genes or enhance mitophagy activity through TBK1 phosphorylation of autophagy adaptors. Upregulation of mitophagy may drive cytosolic DNA accumulation, inflammation, oxidative stress, and mitochondrial dysfunction. UA treatment could break this cycle.

In conclusion, these data show that UA’s senomorphic effect is mediated through targeting NF-κB signaling, as many other senomorphic drugs do, and by targeting the root of the cause SASP – DAMP signaling.

## Methods

### Cell culture and Urolithin A treatment

IMR-90 primary lung fibroblasts (ATCC, CCL-186) were used at population doubling level (PDL) 30-45. Cells were cultured at 37^°^C in an atmosphere with 5% CO2 and 3% O2. IMR-90 cells were cultured in DMEM complete media containing Dulbecco’s Modified Eagle’s Medium (DMEM) (Corning, 10-013-CV) supplemented with 10% (v/v) fetal bovine serum (FBS) (Millipore Sigma, F4135) and penicillin–streptomycin (Corning, 30-001-CI). To confirm the effect of UA on senescent cells, cells were treated 24hrs after seeding (NS) or senescence induction (S) with 20 μM UA in complete growth media. Media was replaced with fresh, containing UA at least 2 times prior to analysis. All cells were mycoplasma-free. Quiescence was induced by replacing culture media with low serum medium (0.2% FBS) 24 hrs before analysis. Cumulative PDL was calculated using the following equation:

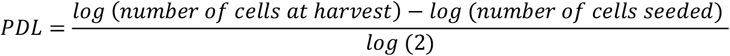

### Primary senescence induction

Senescence was induced by treating cells with 300 nM doxorubicin hydrochloride (Millipore Sigma, 504042) for 24hrs. After 24 hrs, doxorubicin was removed, and the cells were maintained in a complete medium for 9 days. On day 9, the complete medium was changed to a low serum medium for 24 hrs before analysis.

### Conditioned medium (CM) preparation and secondary senescence induction

CM was generated by culturing primary senescent cells in low serum media for 24 hrs before harvesting and counting cells. Collected CM was spun down to remove cell debris, and the supernatant was stored at -80^°^C. All quantitative assays from CM were normalized to the cell number. Secondary senescence was induced by treating proliferating cells 24 hrs after seeding, with 50% CM collected from primary senescent cells and 50% complete media for 5 days. Media was changed on day 5, CM and cells were collected, and multiple senescence markers were tested.

### Senescence-associated beta-galactosidase

SA-β Gal activity was measured using the Senescence Detection Kit (BioVison; K320), following the manufacturer’s instructions. IMR90 cells were plated 1 day before senescence induction in 6-well culture plates (Greiner Bio-One; 657160). Non-senescent cells were plated 3 days before staining. Staining was performed 10 days post-senescence induction. During staining, cells were incubated for 48 hours at 37 ^°^C without CO2 and imaged by brightfield microscopy. The percentage of SA-B-Gal positive cells was counted using ImageJ.

### Immunofluorescence

IMR90 cells per well were plated in 96 well plates and senescence was induced in the plate as described above. Non-senescent cells were seeded 3 days before staining. All immunostaining was performed 10 days after treatment. Cells were fixed with 4% paraformaldehyde in PBS (Thermo Scientific, AAJ19943K2) for 15 minutes at room temperature. After washing with PBS, cells were permeabilized with 0.5% Triton X-100 for 15 minutes, rinsed with PBS, and incubated overnight with primary antibody (1:1000 dilution with 5% BSA) at 4^°^C. Primary antibodies used included γH2AX mouse antibody (Novus Biologicals, NB100-74435) and anti-HMGB1 rabbit antibody (Abcam, ab18256). The following secondary antibodies were used: Alexa Fluor 488 goat anti-mouse (Invitrogen, #A-11008) and Alexa Fluor 546 goat anti-rabbit (Invitrogen, #A-11030). Cells were stained with Hoechst 33342 (Invitrogen, H3570) in 5% BSA for 20 minutes at room temperature in the dark. Cells were washed with PBS before imaging. Cells with >2 γH2AX foci per nucleus and/or the absence of HMGB1 intracellularly were defined as senescent.

Pico green (Thermo Fischer P11496) was combined with the mitochondrial dye – mito-tracker red 633 for extranuclear DNA staining. Both dyes were added to cells in complete media for 1 hour immediately before live cell imaging on a fluorescent microscope. Hoechst staining was used to separate nuclear staining from extranuclear foci.

### Quantitative real-time PCR (Real Time-qPCR)

RNA from cells was extracted using a Quick-RNA Miniprep Kit (Zymo Research, R1055) according to the manufacturer’s instructions. cDNA synthesis was performed using Takara PrimeScript™ RT Master Mix (Takara, RR036A) according to the manufacturer’s instructions. Quantitative PCR was performed on a StepOnePlus™ Real-Time PCR System using primers and probes purchased from Applied Biosystems TaqMan Gene Expression assays. Primers and probes used are listed in Table 1. All results shown as relative to the mean of housekeeping genes (ΔΔCt method), normalizing target cDNA Ct values to that of actin.

**Table 1.**

|  | Thermo Fischer Probe ID |
| --- | --- |
| CDKN1A | Hs00355782_m1 |
| CDKN2A | Hs00923894_m1 |
| IL1A | Hs00174092_m1 |
| IL6 | Hs00174131_m1 |
| IL8 | Hs00174103_m1 |
| CCL5 | Hs00174575_m1 |
| CCL2 | Hs00234140_m1 |

### Enzyme-linked immunosorbent assays (ELISA)

Senescence was induced as indicated above and cultured in low serum medium for 24 hrs before collection. CM was collected and cell debris was removed by centrifugation at 300G for 10 minutes. CM was analyzed with IL-6 and IL-8 ELISA kits (Millipore Sigma) as instructed by the manufacturer and normalized to the cell number.

### Statistical Analysis

Data are presented as mean +/-SEM. All statistical analyses were performed by one-way ANOVA.

## Funding support

We want to acknowledge funding agencies who supported this work by the SENS Research Foundation.

## Conflict of interest

M.C. and J.K.A. are co-founders of Symbiont Bio and declare no financial interests related to this work.

